# Dynamic fluctuations in ascending heart–to–brain communication under mental stress

**DOI:** 10.1101/2022.09.09.507362

**Authors:** Diego Candia-Rivera, Kian Norouzi, Thomas Zoëga Ramsøy, Gaetano Valenza

## Abstract

Dynamical information exchange between central and autonomic nervous systems, as referred to functional brain–heart interplay, occurs during emotional and physical arousal. It is well documented that physical and mental stress lead to sympathetic activation. Nevertheless, the role of autonomic inputs in nervous-system-wise communication under mental stress is yet unknown. In this study, we estimated the causal and bidirectional neural modulations between EEG oscillations and peripheral sympathetic and parasympathetic activities using a recently proposed computational framework for a functional brain–heart interplay assessment, namely the sympathovagal synthetic data generation model. Mental stress was elicited in 37 healthy volunteers by increasing their cognitive demands throughout three tasks associated with increased stress levels. Stress elicitation induced an increased variability in sympathovagal markers, as well as increased variability in the directional brain–heart interplay. The observed heart–to–brain interplay was primarily from sympathetic activity targeting a wide range of EEG oscillations, whereas variability in the efferent direction seemed mainly related to EEG oscillations in the gamma band. These findings extend current knowledge on stress physiology, which mainly referred to top-down neural dynamics. Our results suggest that mental stress may not cause an increase in sympathetic activity exclusively as it initiates a dynamic fluctuation within brain-body networks including bidirectional interactions at a brain–heart level. We conclude that directional brain–heart interplay measurements may provide suitable biomarkers for a quantitative stress assessment and bodily feedback may modulate the perceived stress caused by increased cognitive demand.

## 1. Introduction

Human physiology entails constant and dynamic adaptations in response to cognitive demand through regulatory mechanisms. As part of the regulatory processes, monitoring of peripheral bodily activity contributes to the adaptation to changes in the self or in the environment (1). To illustrate, these processes may stimulate specific behaviors that allow finding shelter or food in extreme conditions. While such physiological adjustments comprehensively refer to “homeostasis”. Adjustments that anticipate future needs refer to allostasis (2). Allostasis thus requires cognitive functions, such as subjective perception, understanding, learning, and memorizing (2).

From a holistic point of view, the physiological responses to cognitive load refer to “mental stress”, which can be elicited by memory, arithmetic, and increased cognitive demand tasks (3). Physical stress involves the physiological responses triggered by homeostatic regulations to bodily conditions, emerging from physical exercise or environmental changes (e.g., temperature or atmospheric pressure) (3). Mental and physical stress encompas physiological responses from different brain structures, together with responses from peripheral systems (4). The neurophysiology of stress sets the hypothalamus as a central component, in which the paraventricular nucleus is the main integrator of stressors, activating systems such as the sympathetic-adreno-medullar and hypothalamus-pituitary-adrenal axes (5). The brain structures actively involved in stress responses include the prefrontal cortex (6) and the amygdala, whose activity is also associated with emotional processing (7). Prefrontal projections to the amygdala (8), as well as hippocampus projections to the amygdala and prefrontal cortex (9) are involved as well. Underlying stress mechanisms have also been captured in EEG studies, showing a high diversity of responses, including hemispheric changes in alpha power and wide-range variability in the EEG spectrum (10, 11).

The central-autonomic network integrates the interoceptive and exteroceptive information to promote physiological and behavioral changes that allow adaption to ongoing challenges, including stress conditions (4, 12–14). Previous studies highlighted a close relationship between stress and sympathetic nervous system activity (15–17), which has also been assessed through series of heart rate variability (18–22), skin conductance (22, 23), breathing (24), body temperature (25, 26) and blood pressure (18, 19, 22). Gastrointestinal (27, 28), endocrine (29), and immune responses (30) were also taken into account to investigate the functional link between stress and sympathetic response. On the other hand, acute stress triggers concurrent fluctuations in heart rate variability and functional connectivity between the central executive and default mode networks (31). Neural responses to heartbeats have been described as a potential indicator of stress, because of the correlations found with sympathetic indexes (22). Similarly with the correlations found between EEG power and autonomic indexes under mental stress (32).

Since stress conditions may induce emotional responses (33), physiological responses to stress (i.e., stress regulation) may be linked to physiological mechanisms of emotion regulation (34). Indeed, while cardiovascular dynamics are modulated by emotions processing (35, 36), modulation activity of the functional brain–heart interactions have been observed under thermal stress and thermoregulatory responses (26, 37), as well as emotional processing (38). Accordingly, cardiac interoceptive feedback seems actively involved under stressful conditions (39, 40), and a wider involvement of the functional brain–peripheral body axis in mental stress has already been hypothesized (41). Nonetheless, the functional brain-peripheral body physiology associated with mental stress is yet unknown. We have hypothesized that the embodiment of mental stress is reflected in bidirectional brain-heart interplay, with specific involvement of sympathetic and vagal dynamics. Accordingly, this study aims to uncover the directional brain– heart interplay mechanisms involved in mental stress induced through visual stimulation and memory tasks. Specifically, we exploited our recently proposed Sympathovagal Synthetic Data Generation model (SV-SDG) (37) to uncover the mutual functional communication between cortical oscillations, as measured through EEG, and cardiac sympathetic/parasympathetic activities, estimated from heartbeat dynamics. The SV-SDG model provides time-varying estimates of the causal interplay between sympathetic/parasympathetic activities and EEG oscillations in a specific frequency band. The framework embeds a heartbeat generation model based on the estimation of sympathetic and parasympathetic activities from Laguerre expansions of the heartbeat series (42).

## 2. Materials and methods

### 2.1 Dataset description

Data were gathered from 37 healthy participants (age median 30 years, age range 22–45 years, 20 males, 17 females) who underwent mental stress elicitation tasks. Participants were asked to sit comfortably and follow instructions on a screen. Recordings of physiological signals included EEG (9-channel, Biopac B-Alert) and one lead ECG, both sampled at 256 Hz.

This study was performed at Neurons Inc, Taastrup, Denmark, in accordance with the Declaration of Helsinki and followed the rules and laws of the Danish Data Protection Agency. Data protection policy also followed the European Union law of the General Data Protection Regulation, as well as the ethical regulations imposed by the Neuromarketing Science and Business Association, Article 6. Each person’s biometric data, survey responses, and other types of data were anonymized and only contained the log number as the unique identifier. Personal information cannot be identified from the log number. The data processing was approved by the ethics committee “Comitato Bioetico di Ateneo” of the University of Pisa

### 2.2 Experimental protocol

Stress induction was performed through a parametric modulation protocol (43, 44) with increased stress level over time. The protocol comprised four stressing conditions, including 1-minute rest and three different stress load tasks, each of which lasted 15 minutes approximately (5 minutes each task). The stressors were presented in the same order to all participants. The first stress load condition consisted in watching a documentary. The second stress load condition consisted in watching a documentary concurrently to performing a digit span task. The third stress load condition consisted in watching a documentary, performing the digit span task and the red box task. For each condition, participants were asked to self-assess and report the perceived stress level through a discrete scale from 1 to 7 (from low to high stress score).

More specifically, the first five minutes of the documentary “The Reality of Van Life”, Different Media © 2018, was projected onto a screen as the first stressor (Figure 1A). The digit span task started with a fixation cross for 1.5 s. Then, three digits were presented for 5 s, followed by a blank screen for 4 s. The participant was then asked to verbally state the three digits in up to 5 s (Figure 1B). The red box task, ran in parallel to the digit span task (Figure 1C), started with a fixation cross for 1.5 s. Then a red box (4×4 red and white box pattern) was presented for 3 s. Next, the three digits were presented for 5 s, followed by a blank screen for 4 s. Then the participant was asked to verbally state the three digits in up to 5 s. Consecutively, a red box was presented, and the participant was asked if the pattern matches to the previously presented one (yes or no answer).

**Figure 1.**
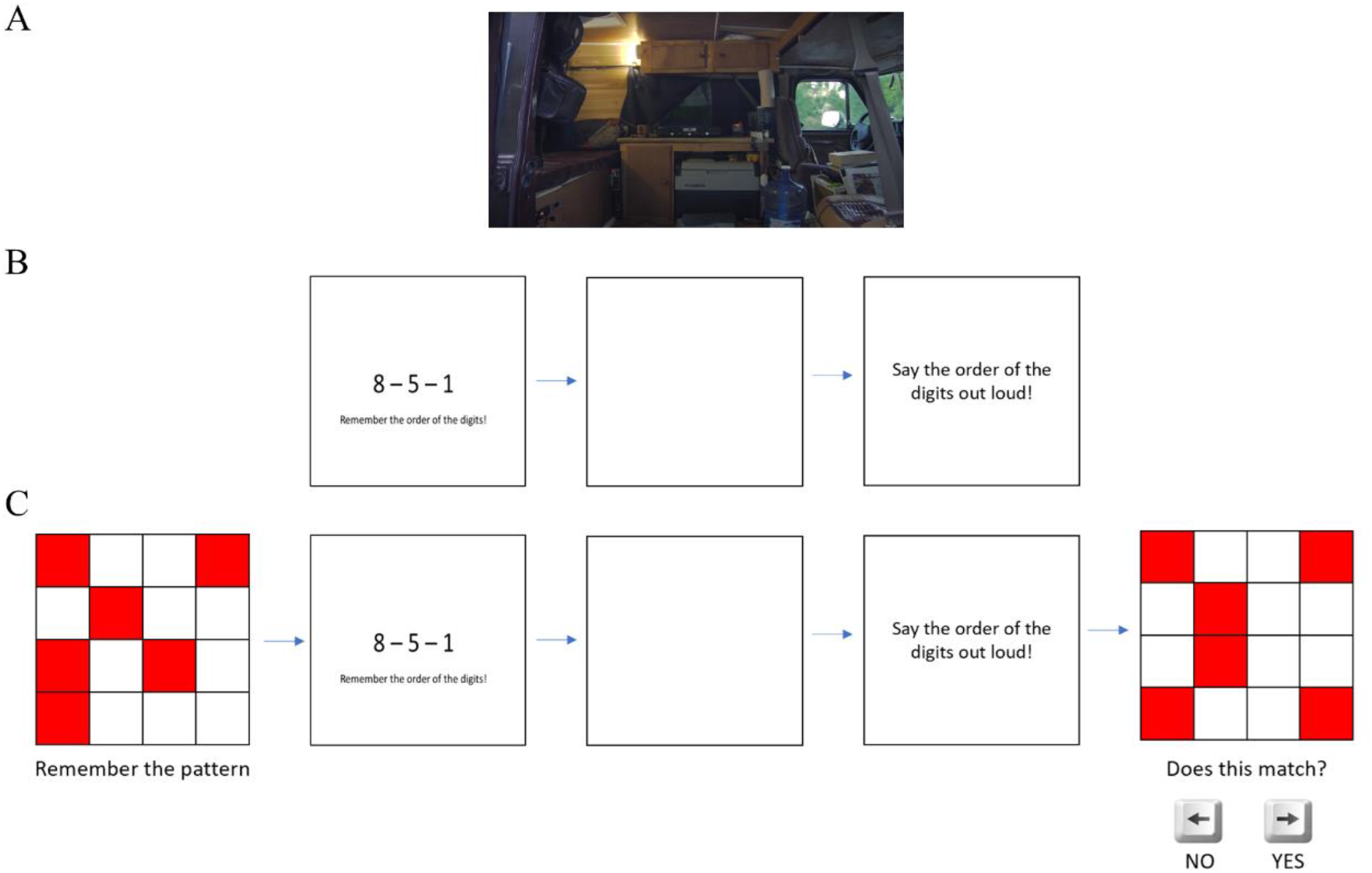
Exemplary stress elicitation images. (A) Sample image from stress load condition 1. The displayed video is an excerpt from the documentary “The Reality of Van Life” (Different Media © 2018). The video was presented for 5 minutes. (B) Sample image from stress load condition 2. The experimental condition consisted in watching the documentary from (A) simultaneously to performing a digit span task of memorizing 3-digit sequences, for approximately 5 minutes. (C) Sample figure from stress load condition 3. The experimental condition consisted in watching the documentary from (A), performing the digit span task from (B), and performing the red box task of memorizing 4×4 patterns, for approximately 5 minutes.

### 2.3 EEG pre-processing

EEG data were pre-processed using MATLAB R2022a and Fieldtrip Toolbox (45). EEG data were bandpass filtered with a Butterworth filter of order 4, between 0.5 and 45 Hz. Large movement artifacts were visually identified and removed manually from independent component space and wavelet filtering. Consecutively, the Independent Component Analysis (ICA) was computed to visually recognize and reject the eye movements and cardiac-field artifacts from the EEG data. One lead ECG was included as an additional input to the ICA to enhance the process of finding cardiac artifacts. Once the ICA components with eye movements and cardiac artifacts were visually identified, they were removed to reconstruct the EEG series. Channels were re-referenced using a common average, which is the most appropriate for a brain–heart interplay estimations (46).

The EEG spectrogram was computed using the short-time Fourier transform with a Hanning taper. Calculations were performed through a sliding time window of 2 seconds with 50% overlap, resulting in a spectrogram resolution of 1 second and 0.5 Hz. Then, time series were integrated within five frequency bands (delta: 1-4 Hz, theta: 4-8 Hz, alpha: 8-12 Hz, beta: 12-30 Hz, gamma: 30-45 Hz).

### 2.4 ECG data processing

ECG time series were bandpass filtered using a Butterworth filter of order 4, between 0.5 and 45 Hz. The R-peaks from the QRS waves were detected in a procedure based on template-matching method (46). All the detected peaks were visually inspected over the original ECG, along with the inter-beat intervals histogram. Manual corrections were performed where needed and guided from the automatic detection of ectopic beats (47).

### 2.5 Estimation of sympathetic and parasympathetic activities

Heart rate variability (HRV) series is usually analyzed by using the Fourier transform, which represents HRV in the frequency domain. This method groups the values into different frequency ranges (VLF: < 0.04 Hz, LF: 0.04-0.15 Hz, and HF: 0.15-0.4 Hz). However, an alternative approach is the use of autoregressive models, which have the advantage of reducing the dimensionality of the frequency-space by defining a limited number of preferred oscillations. This method has been widely used in autonomic assessment, as the frequencies within the HF range can be directly associated with vagal dynamics (48). However, there are limitations to this method, such as the fact that the LF range contains both vagal and sympathetic dynamics (49, 50). To overcome these limitations, we recently proposed the Sympathetic and Parasympathetic Activity Indices (SAI and PAI, respectively), that use Laguerre functions as an alternative way to analyze HRV through autoregressive models. These functions are characterized by a specific “order” and “α” value, which can vary from zero to any positive integer. Indices of sympathetic and parasympathetic activities are derived from characterizing and predicting each heartbeat event by using a combination of past information of RR intervals (R-to-R-peak intervals). This approach improves the identification of model parameters needed to estimate the cardiac sympathetic and parasympathetic activities (48).

Methodologically, the series of RR intervals were convolved with a set of Laguerre functions *φ*_*j*_, as shown in Equation (1):

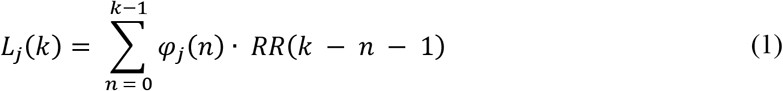

Therefore, the RR series can be expanded using the convolved Laguerre functions *L*(*k*) = [*L*_0_(*k*), *L*_1_(*k*), …, *L*_8_(*k*)]^*T*^, and the theoretical autoregressive model can be used to separate the sympathetic and parasympathetic components as follows:

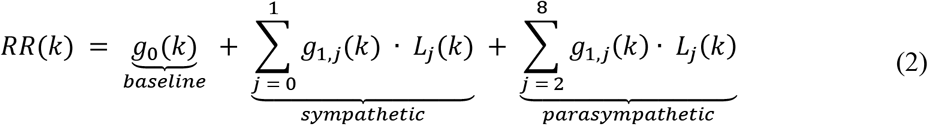

The time-varying Laguerre coefficients *g*(*k*) = [*g*_0_(*k*), *g*_1,0_(*k*), …, *g*_1,8_(*k*)]^*T*^ were modelled according to a dynamic system that fulfils Equations (3) and (4).

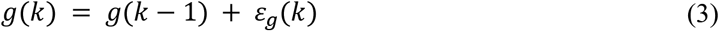

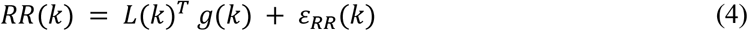

where *ε*_*g*_ is the state noise and *ε*_*RR*_ is the observation noise. The coefficients were then estimated using a Kalman filter with a time-varying observation matrix (51), and SAI and PAI were estimated as shown in Equations (5) and (6).

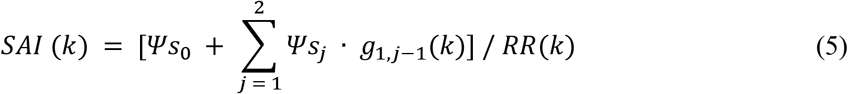

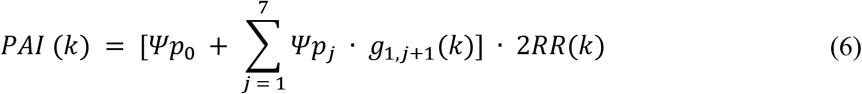

Here, 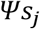 and 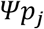 are the generalized values for the sympathetic and parasympathetic kernels with numeric values of 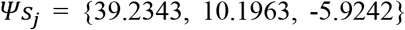 and 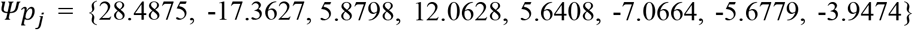. For a comprehensive description of the model generation and parametrization, see (48). SAI and PAI were computed using a publicly available online software, which can be gathered from *www.saipai-hrv.com*.

The validation of SAI and PAI computation has been performed in different studies, including congestive heart failure (52), muscle sympathetic nerve stimulation (53), lower body negative pressure (51), pre-ejection period measurement (54), and controlled breathing (55).

### 2.6 Functional brain–heart interplay assessment

The Sympathovagal Synthetic Data Generation model (SV-SDG) provides time-variant estimates of the bidirectional functional coupling between heartbeat and brain components. The model uses the estimation of sympathetic and parasympathetic activities proposed in (42, 48), described in section 2.5.

#### 2.6.1 Functional Interplay from the brain to the heart

The top-down functional interplay was quantified through a model of synthetic heartbeat generation based on Laguerre expansions of RR series (see Candia-Rivera et al., 2021a for further details). Briefly, heartbeat generation was based on the modulation function *m(t)*, which contains the fluctuations with respect to the baseline heart rate. Such fluctuations were modeled including the sympathetic and parasympathetic interplay. The modulation function *m(t)* was expressed as a linear combination of sympathetic (SAI) and parasympathetic activity index (PAI), and their respective control coefficients *C*_*SAI*_ and *C*_*PAI*_, representing the proportional central nervous system contribution:

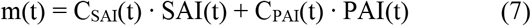

The modulation function was then taken as input to an integrate-and-fire model (42). The model was fitted on the RR interval series using a 15-seconds sliding time window and a linear regression model with no constant term.

C_SAI_ and C_PAI_ coefficients model the directional interaction from EEG activity to the sympathetic and parasympathetic autonomic component, respectively. Accordingly, the directional interaction from cortical oscillations in each band to autonomic component modulating heartbeat dynamics was defined as:

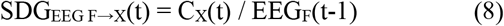

where X ∈ {SAI, PAI}, and *EEG*_F_ indicates the time-varying EEG power with F ∈ {δ, θ, α, β, γ}.

#### 2.6.2 Functional Interplay from the heart to the brain

The functional interplay from heart to brain was quantified through a model based on the generation of synthetic EEG series using an adaptative Markov process (56). The model was fitted using a least-square auto-regressive process to estimate cardiac sympathovagal contributions to the ongoing fluctuations in EEG power as:

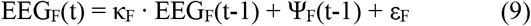

where F is the EEG frequency band, κ_*F*_ is a fitting constant, ε_*F*_ is the adjusted error, and *Ψ*_*F*_ indicates the fluctuations of EEG power in *F*. Then, the heart–to–brain functional coupling coefficients were calculated as follows:

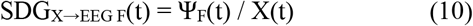

where X ∈ {SAI, PAI}. For further details, please see Candia-Rivera et al., 2022a.

The software for computation of SAI and PAI is available at *www.saipai-hrv.com*. The source code implementing the SV-SDG model is available at www.github.com/diegocandiar/brain_heart_svsdg.

### 2.7 Multivariate analysis

In order to identify the most significant brain-heart features sensitive to mental stress, a multivariate analysis was performed. The feature selection is based on the ranking provided by the computation of Minimum Redundancy Maximum Relevance (MRMR) scores (57) and was computed over the 180 SV-SDG-derived features (180 = 2 directions x 2 autonomic markers x 5 brain oscillations x 9 channels) to select the five most significant ones in two conditions: (i) a linear regression model predicting the median stress level in each condition, and (ii) a binary classification algorithm to discern low vs high stress level.

The MRMR score computation algorithm was performed as follows:

1. The relevance *V*_*x*_ of all features *x* was computed. The feature with the largest relevance 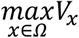 was selected. The selected feature was added to an empty set of features *S*. *V*_*x*_ was defined as:

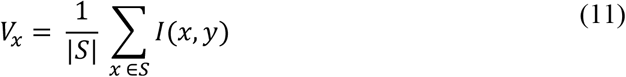

Where *S* ∨ is the number of features in *S* and *I*(*x, y)* is the mutual information between the feature x and the output y:

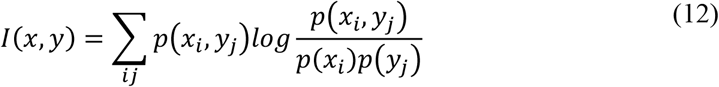
2. Next, the features with non-zero relevance *V*_*x*_ and zero redundancy *W*_*x*_ in *S*^*c*^ (complement of *S*) were identified. Then, the feature with the largest relevance was selected, 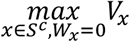. The selected feature was added to the set *S*. *W*_*x*_ was defined as:

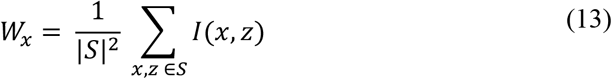

If *S*^*c*^ did not include a feature with non-zero relevance and zero redundancy, skip step number 3
3. Step number 2 was repeated until the redundancy *W*_*x*_ was not zero for all features in *S*^*c*^.
4. The feature with the largest *MIQ* was selected, with non-zero relevance and non-zero redundancy in *S*^*c*^, and the selected feature was added to the set *S*. *MIQ* was defined as:

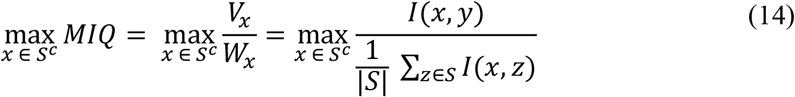
5. Step 4 was repeated until the relevance was zero for all features in *S*^*c*^.
6. The features with zero relevance were added to *S* in random order.

The multivariate analyses were performed in a 5-fold cross-validation framework. Linear regressions to the stress level were performed using least squares kernel regression with regularization strength set to 0.027. The stress level was quantified “0” at rest, “1” for stressor 1, “4” for stressor 2, and “5” for stressor 3 to closely match the median stress ratings from subjects’ self-assessment reports. The regression performance was measured through Root Mean Squared Error (RMSE) for the prediction of median stress ratings. Binary classification for the low vs. high stress recognition was performed through a kernel naïve Bayes classifier with a Gaussian kernel, with “low stress” class associated with “rest” and “stressor 1” conditions, and “high stress” associated with the stressors 2 and 3. The classification performance was quantified through the classification accuracy.

### 2.8 Statistical analysis

Group-wise statistical analysis between resting state and the three stressor levels was performed through non-parametric Friedman tests, whereas two-condition comparisons were performed through Wilcoxon signed-rank test. The statistical testing was performed per EEG channel, in which the inputs correspond to SV-SDG coupling coefficient computed at different experimental conditions. The significance level of the *p*-values was corrected in accordance with the Bonferroni rule for 9 channels, with an uncorrected statistical significance set to alpha = 0.05. The samples were described group-wise using the median and related dispersion (variability) measures that was quantified though the median absolute deviation (MAD).

## 3 Results

The participants’ self-reports on the perceived level of stress are displayed in Figure 2 for each stressful condition, where the group median ± MAD reported stress levels are 1±0, 4±1 and 5±1 (p = 2 · 10^−14^ from Friedman test). A multiple comparison analysis showed that the three stressful conditions are significantly different (p<0.00005).

**Figure 2.**
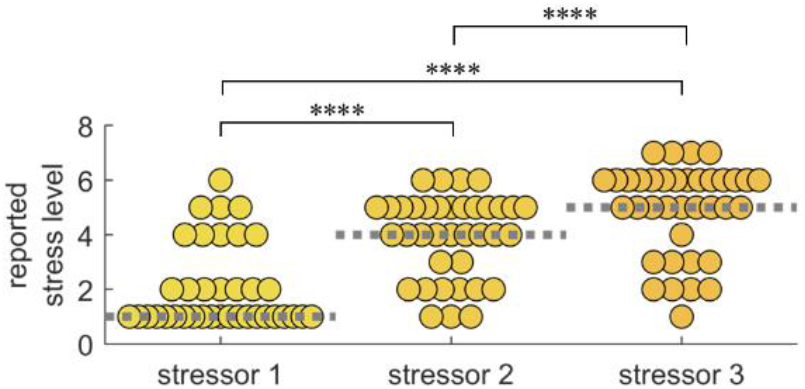
Self-reported stress level for three stressful conditions. Each data point corresponds to the reported stress level per subject for each for: i) stress load condition 1: documentary, ii) stress load condition 2: documentary + digit span task, iii) stress load condition 3: documentary + digit span task + red box task. **** < 0.00005 from Wilcoxon signed-rank test.

Cardiac autonomic activity was assessed through the sympathetic and parasympathetic activity indices (SAI and PAI, respectively). While condensing the SAI and PAI time-resolved information, median SAI and median PAI did not change significantly across the experimental conditions (p = 0.0935 from Friedman test on median SAI, and p = 0.3101 from Friedman test on median PAI). Nevertheless, SAI and PAI variability (i.e., MAD over time) significantly changes across the experimental conditions (p = 7 · 10^−6^ from Friedman test on SAI variability and p = 4 · 10^−9^ from Friedman test on PAI variability). Figure 3 depicts group-wise distributions for RR, SAI and PAI median and variability, with evident increase in SAI and PAI variability in the three stressful conditions as compared to rest.

**Figure 3.**
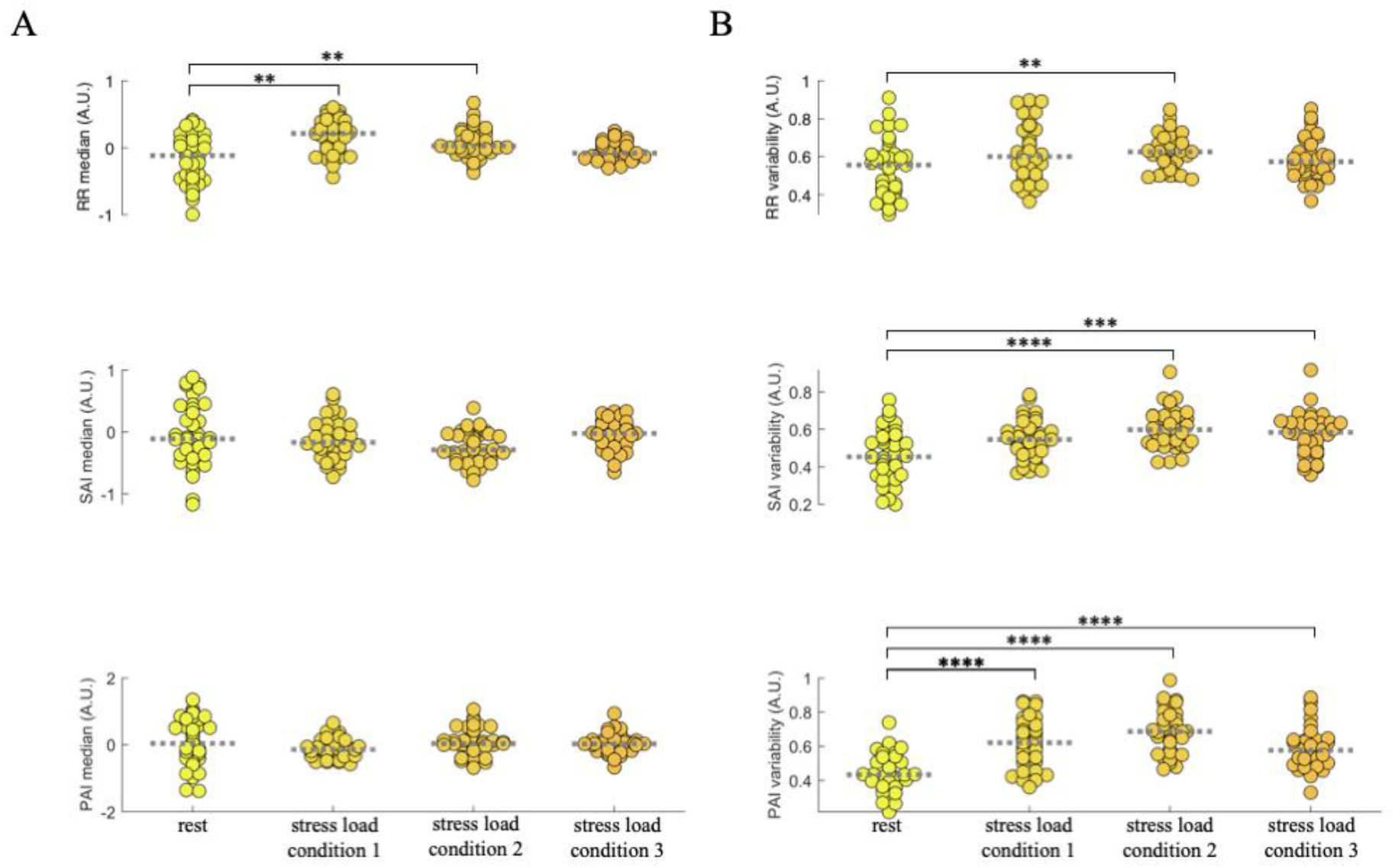
Group-wise distributions of RR, SAI and PAI median and variability for each experimental condition. Each data point corresponds to the measured autonomic marker per subject at each of the four conditions. (A) RR, SAI and PAI median. (B) RR, SAI and PAI variability as measured through median absolute deviation (M.A.D.). The time-varying autonomic indexes were z-score normalized for the whole experimental protocol duration before computing median and M.A.D values. ** p < 0.005, *** p < 0.0005, **** p < 0.00005 (Bonferroni-corrected significance at α < 0.00833).

Since autonomic variability is sensitive to stress levels, we further explored how they relate to brain–heart interplay. Figure 4 illustrates results from the Friedman tests on group-wise brain-heart variability changes among experimental conditions. Most of the significant changes among conditions are associated with ascending interactions, especially originating from sympathetic and vagal activity targeting EEG oscillations in the alpha band (SAI→alpha: all channels Friedman test p ≤ 0.00013, PAI→alpha: all channels Friedman test p ≤ 0.00003). Ascending heart-to-brain communication targeting EEG oscillations in the theta, beta and gamma bands show significant changes as well, together with descending interactions from cortical gamma oscillations to vagal activity (see Table 1). In contrast, cortical power variability mostly shows not significant changes, with a few statistical differences associated with gamma oscillations in the left-frontal electrodes, as shown in Table 2 (see Supplementary Fig. 2 for visualizing an exemplary subject’s SAI, PAI and gamma power fluctuations on time).

**Figure 4.**
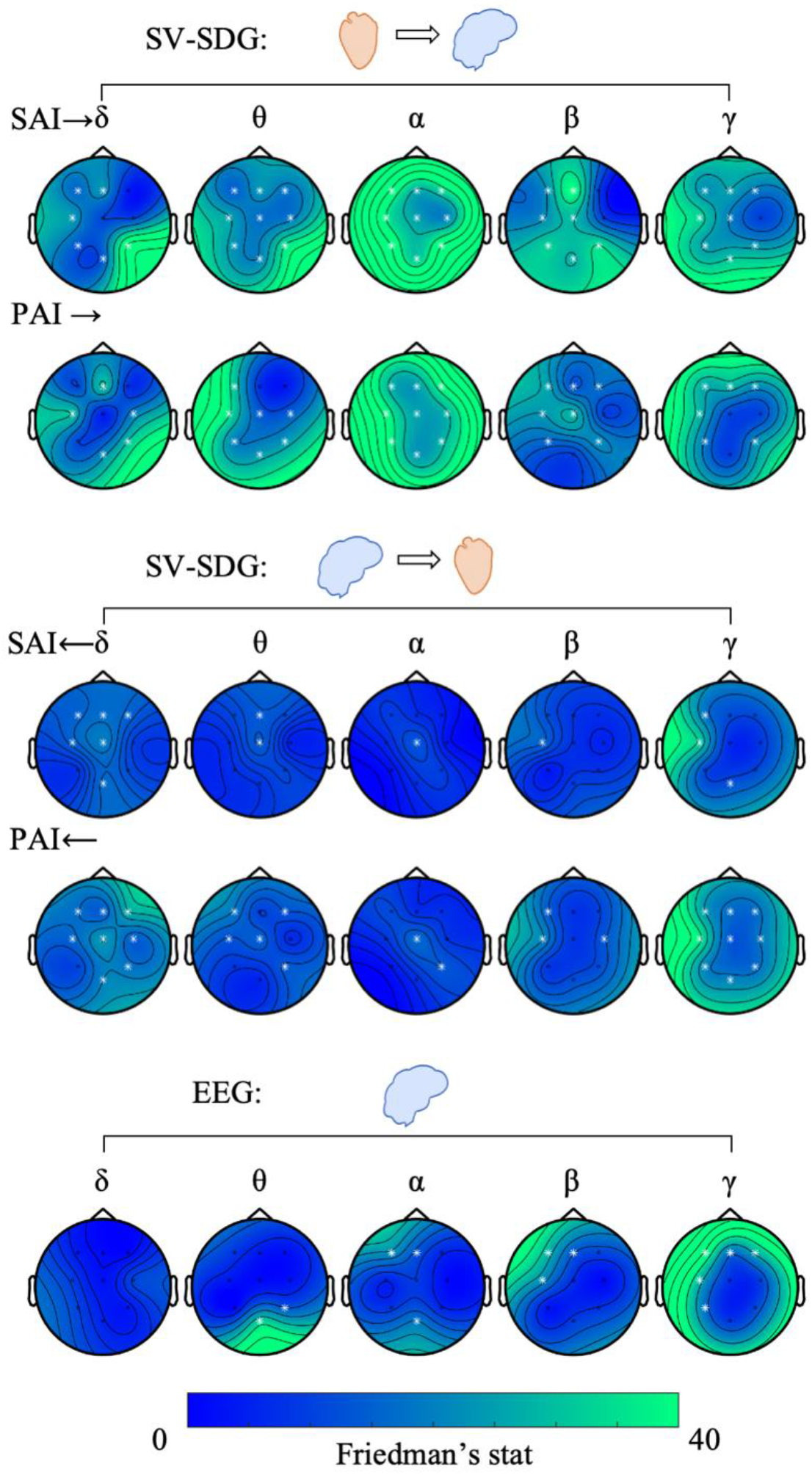
Topographic maps of Friedman non-parametric test for paired brain–heart interplay variability on the ascending (top panel) and descending (middle panel) direction from the SV-SDG model among experimental conditions (rest and three mental stress conditions). In the bottom panel, topographic maps of Friedman non-parametric test for paired EEG power variability among experimental conditions (rest and three mental stress conditions) are shown. White electrodes indicate p < 0.0056. The variability measure was the median absolute deviation. SV-SDG: Sympathovagal synthetic data generation, SAI: Sympathetic activity index, PAI: Parasympathetic activity index. Complementary results on SV-SDG and EEG power medians are in Supplementary Figure 1.

**Table 1.** Friedman test results on the variability of brain-heart interplay coupling coefficients. Median p-values and Z-values among significant channels are displayed. Critical alpha was set according to the Bonferroni rule for multiple comparisons among channels at α = 0.05/9 ≈ 0.0056. Variability of coupling coefficients on time was computed with median absolute deviation (MAD).

| Ascending coupling coefficients | Median p-value (among significant channels) | Median Friedman's stat (among significant channels) | Significant channels | Descending coupling coefficients | Median p-value (among significant channels) | Median Friedman's stat (among significant channels) | Significant channels |
| --- | --- | --- | --- | --- | --- | --- | --- |
| SAI→delta | 0.0006 | 19.5789 | F3, Fz, C3, P3, POz, P4 | delta→SAI | 0.0013 | 15.7895 | F3, Fz, F4, C3, Cz, POz |
| SAI→theta | 0.0001 | 20.7789 | F3, Fz, F4, C3, Cz, C4, P3, POz, P4 | theta→SAI | 0.0030 | 13.9579 | Fz, Cz, |
| SAI→alpha | < 0.0001 | 33.8211 | F3, Fz, F4, C3, Cz, C4, P3, POz, P4 | alpha→SAI | 0.0046 | 13.0105 | Cz |
| SAI→beta | < 0.0001 | 28.7053 | F3, Fz, C3, Cz, P3, POz, P4 | beta→SAI | 0.0019 | 14.9368 | C3 |
| SAI→gamma | < 0.0001 | 24.0632 | F3, Fz, F4, C3, Cz, P3, POz, P4 | gamma→SAI | 0.0004 | 18 | F3, C3, POz |
| PAI→delta | < 0.0001 | 28.9263 | Fz, C3, C4, POz, P4 | delta→PAI | 0.0006 | 17.5737 | F3, Fz, F4, C3, Cz, C4, POz, P4 |
| PAI→theta | < 0.0001 | 23.6211 | F3, C3, Cz, C4, P3, POz, P4 | theta→PAI | 0.0033 | 13.7368 | F3, F4, C3, Cz, P4 |
| PAI→alpha | < 0.0001 | 27.6947 | F3, Fz, F4, C3, Cz, C4, P3, POz, P4 | alpha→PAI | 0.0022 | 14.9368 | Cz, P4 |
| PAI→beta | < 0.0001 | 21.6947 | F3, Fz, F4, C3, Cz, P3, P4 | beta→PAI | 0.0025 | 14.3053 | F3, C3, C4 |
| PAI→gamma | < 0.0001 | 24.4421 | F3, Fz, F4, C3 P3, P4 | gamma→PAI | 0.0003 | 18.7579 | F3, Fz, F4, C3, Cz, C4, P3, POz, P4 |

**Table 2.** Friedman test results on the variability of EEG power. Median p-values and Z-values among significant channels are displayed. Critical alpha was set according to the Bonferroni rule for multiple comparisons among channels at α = 0.05/9 ≈ 0.0056. Variability of coupling coefficients on time was computed with median absolute deviation (MAD).

| EEG power | Median p-value (among significant channels) | Median Friedman's stat (among significant channels) | Significant channels |
| --- | --- | --- | --- |
| delta | - | - | - |
| theta | 0.0021 | 22.8158 | POz, P4 |
| alpha | 0.0003 | 18.7895 | F3, Fz, POz |
| beta | < 0.0001 | 21.7579 | F3, Fz, C3 |
| gamma | < 0.0001 | 26.8105 | F3, Fz, F4, C3, P3 |

For the sake of completeness, results on the median brain-heart are shown in Supplementary Figure 1. Mental stress mainly modulates heart-to-brain functional communication, especially targeting delta, alpha, beta (in the left hemisphere), and gamma bands.

According to the MRMR algorithm, the five most informative features for the linear regression analysis and the low vs. high stress classification are reported in Table 3 and depicted in Fig. 5. In both multivariate analyses, ascending features from SAI and PAI are prevalent. To illustrate, while median stress level prediction mostly uses SAI→beta, most of the information needed for low vs high stress classification was provided by PAI→gamma.

**Table 3.** Brain-heart interplay feature ranking according to the Minimum redundancy maximum relevance (MRMR) algorithm. The MRMR scores were computed for two models: regression to the group-median reported stress (rest=0, stress condition 1=1, stress condition 2=4, stress condition 3=5), and classification of low and high stress levels (low=rest and stress condition 1, high=stress conditions 2 and 3).

|  | 1<sup>st</sup> feature | 2<sup>nd</sup> feature | 3<sup>rd</sup> feature | 4<sup>th</sup> feature | 5<sup>th</sup> feature |
| --- | --- | --- | --- | --- | --- |
| Regression | SAI→$\beta$, Cz<br>MRMR=0.2395 | PAI→$\alpha$, F3<br>MRMR=0.2247 | PAI→$\gamma$, F3<br>MRMR=0.1771 | SAI→$\theta$, C3<br>MRMR=0.1669 | PAI→$\delta$, P4<br>MRMR=0.1521 |
| Binary classification | PAI→$\gamma$, C3<br>MRMR=0.1464 | SAI→$\beta$, P3<br>MRMR=0.1399 | SAI→$\delta$, P4<br>MRMR=0.1156 | PAI→$\beta$, P4<br>MRMR=0.0473 | SAI→$\delta$, F3<br>MRMR=0.0453 |

**Figure 5.**
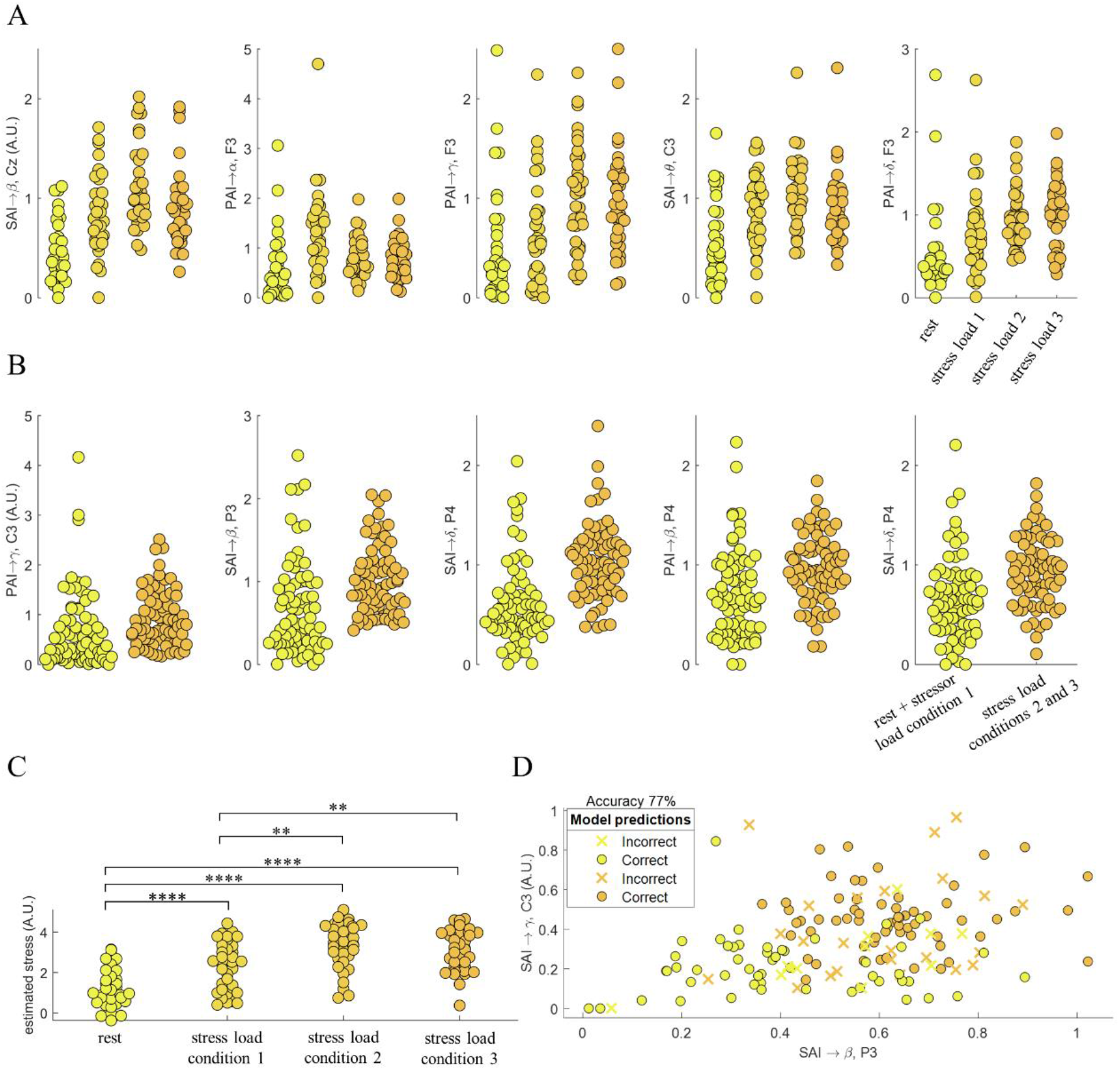
Multivariate analysis results. (A) The five most informative brain–heart interplay features according to the MRMR regression criteria. The regression was performed on the group median stress level: rest=0, stressor condition 1=1, stressor condition 2=4, stressor condition 3=5. (B) The five most informative brain–heart interplay features according to the MRMR criteria to classify low vs high stress levels. Low stress referred to rest and stress load condition 1; high stress referred to stress load conditions 2 and 3. (C) Regression model output from a 5-fold cross-validation to the stress level using the best five markers presented in (A). The statistical comparisons correspond to paired Wilcoxon tests on the estimated stress level score computed with the regression model. (D) Binary stress level classification using the five most informative features in (B). Yellow and orange circles indicate low and high stress conditions, respectively. ** p < 0.005, *** p < 0.0005, **** p < 0.00005 (Bonferroni-corrected significance at α < 0.00833).

In the regression analysis, the RMSE was 1.6851, and its output showed a significant difference between all predicted stress levels but stressor 2 vs stressor 3 (Figure 5C). In the classification, the five brain-heart features achieved a discrimination accuracy as high as 77% (Figure 5D), with a sensitivity of 85.14% on detecting high stress, and 68.92% specificity.

Figure 6 shows exemplary SAI→beta and SAI→delta estimates from one subject for the whole duration of the experimental protocol. An overall increased variability of both markers can be observed in stressful conditions 2 and 3 with respect to rest and stressful condition 1.

**Figure 6.**
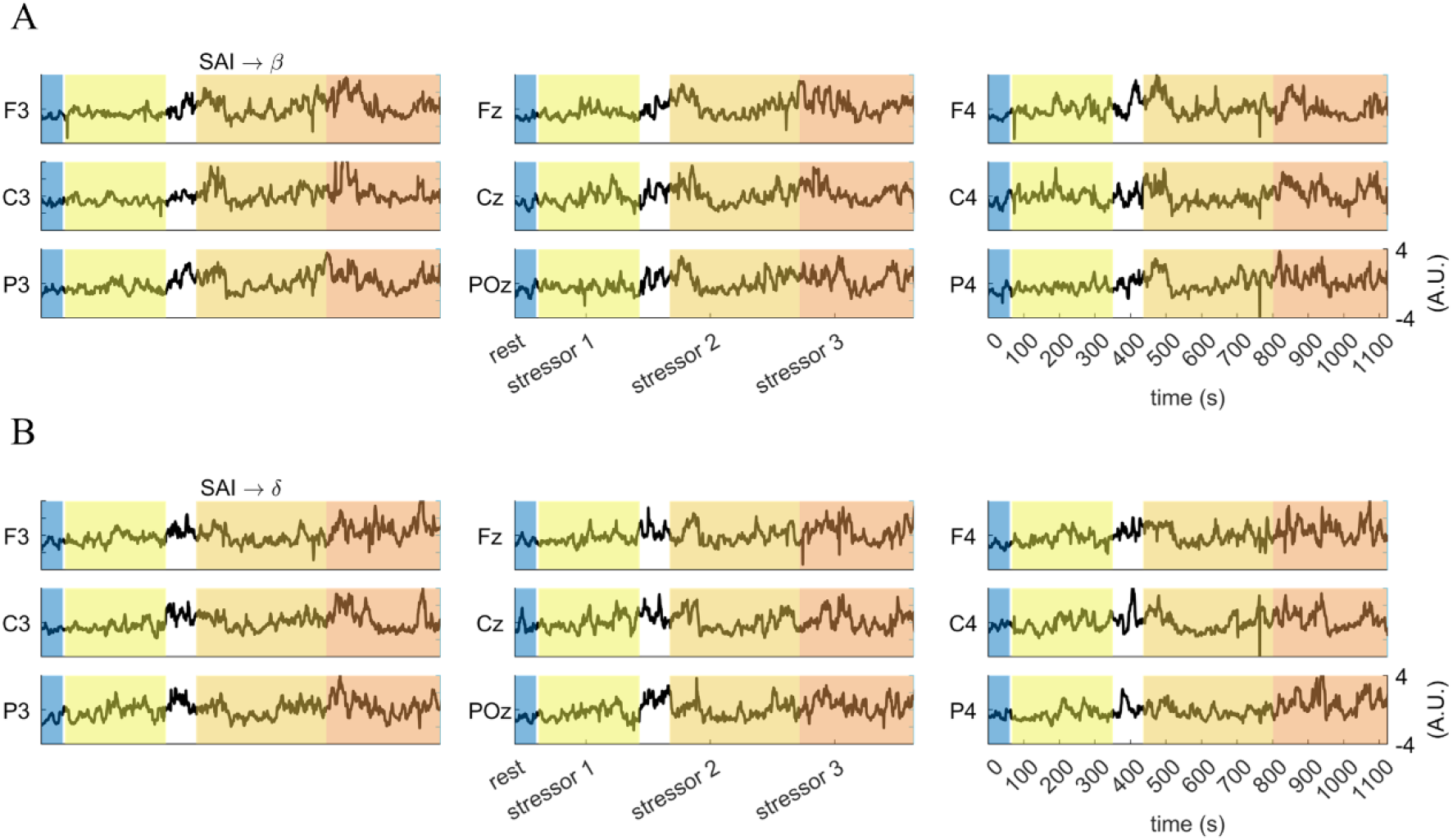
Exemplary participant during the experimental protocol. Parallel fluctuations in (A) SAI→beta and (B) SAI→delta are displayed.

## 4 Discussion

Supported by previous evidence linking mental stress with sympathetic activity (15–17) and emotional responses, we investigated functional brain–heart interplay directionally under hypothesis of modulation across different stress levels.

When condensing the temporal dynamics of sympathetic and parasympathetic activities throughout the experimental conditions, on the one hand, we observed that SAI and PAI central tendencies (median) did not change among stress levels. On the other hand, we observed that the variability (MAD) of SAI and PAI significantly increased in accordance with stress levels up to stressful condition 2. Sympathetic activity, as measured through systolic blood pressure, heart rate, ventricular ejection fraction, and skin conductance has been associated with mental stress (22); moreover, mental stress induced by mental arithmetic increases heart rate variability power in the low frequency and a decrease in its high-frequency power (58–61), suggesting an increase in the sympathetic tone, and a decrease in the parasympathetic one. Stress also modulates heartbeat non-linear dynamics (20, 62). Changes in attention have been referred as a source of autonomic variability (63). Furthermore, some studies have suggested that high-frequency fluctuations in heartbeat dynamics are associated with memory retrieval, reaction time, and action execution (59, 64, 65), suggesting a dynamic interaction between sympathetic and parasympathetic activities under stress elicitation. As stress elicitation may involve some executive functions (e.g., self-control and working memory), the role of high-frequency autonomic activity has been associated with specific dimensions of executive functioning (66–69).

We observed differences among stressful conditions in EEG oscillations in the gamma band. The existing evidence on EEG and stress shows heterogeneous and divergent findings with respect to frequency bands. To illustrate, some studies suggest that different dimensions of stress are associated with alpha-beta interactions (11, 60, 70), theta-beta interactions (70, 71), alpha-gamma interactions (72–74), and theta-alpha interactions (73). Such heterogeneity may be related to the subjectivity and thus high inter-subject variability on perceived stress (72), as well as to the coping strategies (75). For instance, the processing of concurrent inputs/tasks requires multiple access to the working memory (76). Another source of variability may be associated with the level of cognitive demand of the tasks, which may not be directly related to the stress level (77). In this study, the digit span task involved a verbal report, which is also associated with EEG signatures of parieto-occipital desynchronizations of lower alpha (78).

We observed EEG activity modulation as linked to both autonomic branches, whose activity was measured through SAI and PAI. Particularly, our results on the brain–heart interplay shows that the variability of ascending heart–to–brain communication reflects the level of stress, especially until stressful condition 2, as compared to descending brain–to–heart modulations. Indeed, the highest stress level is not statistically associated with the highest variability of heart– to–brain modulation, nor with SAI and PAI dynamics. We speculate this may be due to the following main factors: (i) perceived stress level is mitigated or masked by mental fatigue due to sustained attention (79); (ii) the increasing stress conditions may be subject to an attentional-bradycardic effect to hyper-arousing conditions (80, 81), also known as “freezing” effect, and thus highest stress conditions may be associated with a different physiological response than other stressful conditions.

Previous studies on physiological correlates of stress focused on top-down mechanisms exclusively (7, 82). While brain responses may precede cardiac responses, as measured through EEG (60, 74) and fMRI (83, 84), stressors may elicit activity in the amygdala and hippocampus such that a subsequent bottom-up control is activated (85). Indeed, the brain and heart continuously influence each other (84), and the ascending arousal system shapes brain dynamics to mediate awareness of mental states (86), as well as to facilitate performance at different tasks (87, 88) and to shape physical and emotional arousal (37, 38). Stress regulation shares mechanisms involved in emotion regulation as well (34). To illustrate, the anterior insula integrates interoceptive signals during emotional and cognitive processing, being these processes involved in the monitoring of the physiological state of the body (89). The neural monitoring of cardiac inputs may trigger physiological adjustments in the frame of homeostatic and allostatic regulations under emotion elicitation (36, 90). The functional brain–heart interplay under stress elicitation has been shown in heartbeat-evoked potentials correlating with stress-induced changes in cardiac output (22), and correlates of functional connectivity with heart rate variability (31). The role of cardiac inputs in the neurophysiology of stress is also supported by the experimental evidence showing an increased information flow from heart-to-brain during increased attention (63) and disrupted abilities on detecting cardiac and respiratory signals from oneself under anxiety (91, 92).

On the bottom-up modulation, we observed that both sympathetic and vagal oscillations map onto various EEG oscillations at different frequency bands. Indeed, the sympathetic origin of brain-heart interplay in stress was expected because of previous evidence (18, 19, 22, 23). In this study, SAI→beta interplay seems more sensitive to changes in stress levels. The involvement of beta waves in mental stress has been previously reported (93), along with alpha-beta interactions (11, 60, 70) and theta-beta interactions (70, 71). Note that EEG oscillations in the theta band have been consistently reported as a sensitive correlate of emotion processing (94) and also in a heart–to–brain communication (38). Our results show that preferential heart–to–brain communication occur over the frontal and parietal cortical regions, consistently with a previous report on stress (6) and correlates of cognitive operations (95).

We showed that a multivariate analysis helps to distinguish between stress levels, as compared to individual autonomic markers. While the use of low-density EEG in this study is certainly a limitation to understanding the brain mapping and cortical dynamics of stress neurophysiology, it proves the suitability of this kind of device to detect levels of stress with potential commercial applications. The study of mental stress elicited in other paradigms, such as mental arithmetic, could give a broader view of the physiological processes involved in brain– heart information exchange. Our study confirms the advantages of analyzing the interactions between the brain and heart, instead of studying heart rate and brain dynamics exclusively (96, 97). The understanding of brain–heart dynamics and the neurophysiological substrate of stress has clinical relevance. Heart rate variability markers are acknowledged to reflect autonomic dysregulation, which may lead to morbidity and mortality (98, 99). The evidence also shows differences in heart rate variability between healthy humans and different mood disorders, but also as a marker of the effects of antidepressant medications (98). The description of stress mechanisms can enlighten the apparent relationships with cardiac death (100), cardiovascular disease (101), sudden death (102), and psychiatric disorders (103). The evidence in other markers of brain–heart interplay shows as well that the dynamic interaction of these systems may relate to different aspects of mental health (22, 104, 105).

This study comes with limitations. The self-assessment measurements we used for subjective stress evaluation do not refer to a specific psychometric test. Moreover, we are aware that confounding factors and artifact sources are numerous in EEG and therefore in functional brain-heart interplay studies (106–108)., In particular, the gamma activity is quite sensitive to muscle artifacts. Since simultaneously electromyography recordings were not available, we cannot exclude that all of the artifacts have been rejected in our pre-processing stage. This indeed constitutes a limitation of our study. However, by also relying on the several pre-processing steps and by checking all series by visual inspection, we are confident that our results are reliable in highlighting a functional, directional link between EEG oscillations in the gamma band and sympathovagal oscillations.

## 5 Conclusions

Stress neurophysiology involves bidirectional interactions between the brain and heart, with peripheral bodily feedback playing a key role. Mental stress leads to increased variability in sympathetic and parasympathetic activities, which is also reflected in changes in EEG gamma activity. These results are in line with the experimental evidence showing a dynamic information exchange between the central and autonomic nervous systems during emotional arousal and physical stress. Estimates of functional brain–heart interplay may be suitable biomarkers of mental stress.

## 6 Acknowledgements

The research leading to these results has received funding from the European Commission - Horizon 2020 Program under grant agreement n° 813234 of the project “RHUMBO”.

## 7 Authors’ contributions

Conceptualization: D. Candia-Rivera and T. Z. Ramsøy; Data curation: K. Norouzi; Formal analysis and Investigation: D. Candia-Rivera and G. Valenza; Methodology: D. Candia-Rivera; Supervision: T. Z. Ramsøy and G. Valenza; Writing - original draft: D. Candia-Rivera; Writing - review & editing: All authors.

