## Supplementary material for "Dynamic fluctuations in ascending heart–to–brain communication under mental stress"

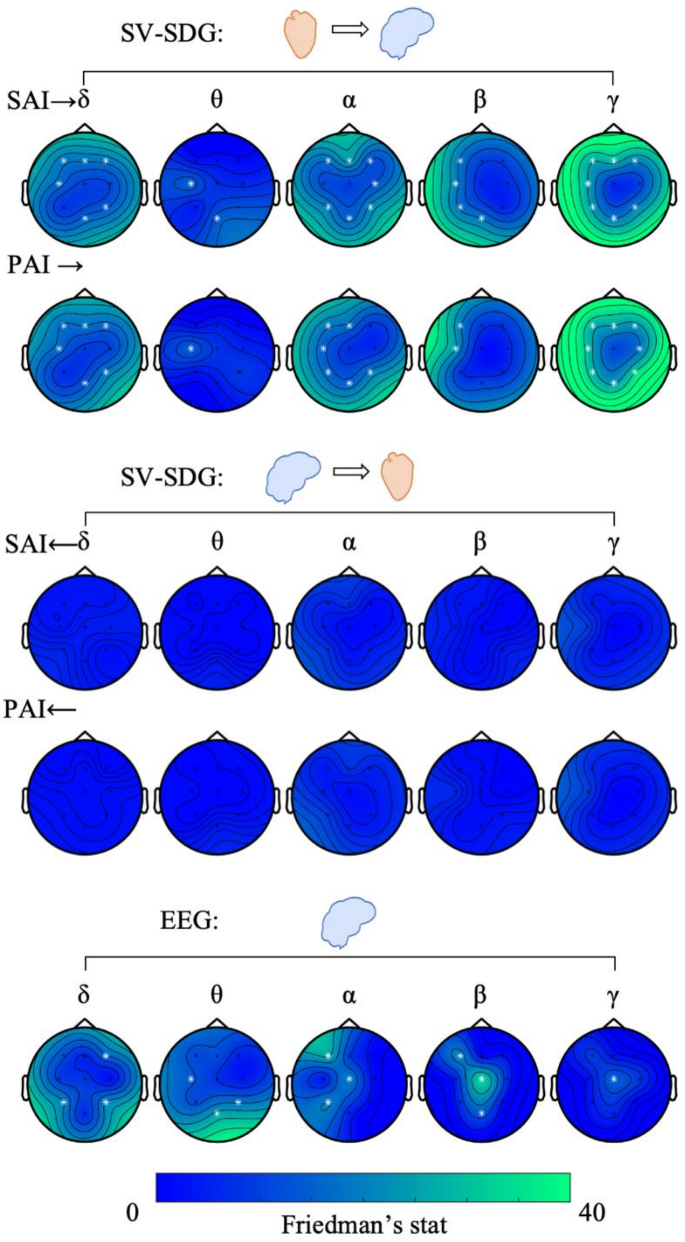


*Supplementary Figure 1. Non-parametric analysis of variance of brain–heart interplay medians (SV-SDG, ascending and descending) and EEG power variability across experimental conditions. Friedman tests were performed at the four experimental conditions (rest and three mental stress conditions). Colormaps indicate the Friedman test statistic. White electrodes indicate p < 0.0056. SV-SDG: Sympatho-vagal synthetic data generation, SAI: Sympathetic activity index, PAI: Parasympathetic activity index.*

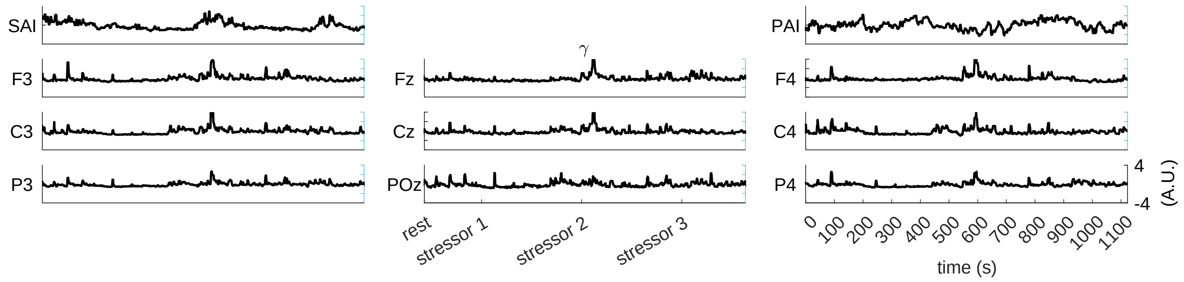


*Supplementary Figure 2. Exemplary participant during the experimental protocol. Parallel fluctuations in SAI, PAI and gamma power are displayed.*
